# Elevation of major constitutive heat shock proteins is heat shock factor independent and essential for establishment and growth of Lgl loss and Yorkie gain mediated tumors in *Drosophila*

**DOI:** 10.1101/2021.08.30.458144

**Authors:** Gunjan Singh, Saptomee Chakraborty, Subhash C. Lakhotia

## Abstract

Cancer cells generally overexpress heat shock proteins (Hsps), the major components of cellular stress response, to overcome and survive the diverse stresses. However, the specific roles of Hsps in initiation and establishment of cancers remain unclear. Using loss of Lgl-mediated epithelial tumorigenesis in *Drosophila,* we induced tumorigenic somatic clones of different genetic backgrounds to examine the temporal and spatial expression and roles of major heat shock proteins in tumor growth. The constitutively expressed Hsp83, Hsc70 (heat shock cognate), Hsp60 and Hsp27 show elevated levels in all cells of the tumorigenic clone since early stages, which persists till their transformation. However, the stress-inducible Hsp70 is expressd only in a few cells at later stage of established tumorous clones that show high F-actin aggregation. Intriguingly, levels of heat shock factor (HSF), the master regulator of Hsps, remain unaltered in these tumorous cells and its down-regulation does not affect tumorigenic growth of *lgl*^−^ clones overexpressing Yorkie, although down-regulation of Hsp83 prevents their survival and growth. Interestingly, overexpression of HSF or Hsp83 in *lgl*^−^ cells makes them competitively successfully in establishing tumorous clones. These results show that the major constitutively expressed Hsps, but not the stress-inducible Hsp70, are involved in early as well as late stages of epithelial tumors and their elevated expression *in lgl*^−^ clones co-overexpressing Yorkie is independent of HSF.

## Introduction

Malignant transformation, which normally remains protected by multiple tumor suppressing mechanisms, involves drastic changes in phenotypes of a di□erentiated cell (Hanahan and Weinberg, 2000; 2011). In epithelial carcinomas, the transformed cells proliferate abnormally despite lack of growth stimuli, develop resistance to apoptosis and senescence, show de novo angiogenesis, and invade surrounding tissues leading to metastasis and colonization of distant tissues (Hanahan and Weinberg, 2011). Since these properties are forbidden for di□erentiated cells, malignant transformation requires several sets of changes in regulatory circuits, facilitated by mutations and reprogramming of cell metabolism (Vogelstein et al., 1988; Yokota, 2000). Such changes expose the cancer cells to a variety of stresses like hypoxia, lack of nutrients, DNA damage, immune responses etc (Calderwood and Murshid, 2017; Calderwood and Gong, 2016; Hanahan and Weinberg, 2011; Lang et al., 2019). Levels of different heat shock protein (Hsp) family members and their master regulator, the heat shock transcription factor (HSF), are elevated in a remarkably wide range of tumor cells, which indicates poor prognosis in most cancer types (Calderwood and Murshid, 2017; Meng et al., 2010). However, their specific roles in initiation and establishment of cancers are still incompletely understood.

The Hsps are a group of molecular chaperones that are highly induced during proteotoxic and other cell stress conditions in HSF dependent manner (Richter et al., 2010; Lindquist and Craig, 1988). and maintain the structural integrity of the proteome to promote proteostasis and cell viability (Calderwood, 2007; Arya et al., 2007). Most Hsps are constitutively present in a cell while a few are completely absent or weakly expressed in unstressed cells but get turned on at dramatic high rates following stress (Feder and Hofmann, 1999; Arya et al., 2007). The significance of elevated Hsps in cancer cells in mediating several hallmarks of cancer is not yet fully understood (Calderwood and Gong, 2016; Calderwood and Murshid, 2017; Lang et al., 2019; Zaarur et al., 2006) Therefore, a good understanding of the cross talks between heat shock response and other signalling pathways is necessary for a deeper understanding of tumorigenesis. In view of the high conservation of pathways known to be involved in tumorigenesis between mammals and flies (Ugur et al., 2016; Millburn et al., 2016; Mirzoyan et al., 2019; Gangwani et al., 2020), and the availability of enormous genetic resources and versatile toolkits like directed gene expression, transgenic and gene-knockout strategies, *Drosophila* offers convenient model system for modulation of the various signaling pathways controlling cell growth and invasion.

There has been no report on expression and roles of Hsps during different stages tumor growth in *Drosophila.* Therefore, we have standardised a fly model for a systematic temporal mapping of expression of different cell stress response genes during the tumor growth. We primarily utilized tumors induced in larval wing imaginal discs using the *l(2)gl^4^ (lgl^−^)* null mutant allele which results in loss of function of the cell polarity Lgl protein. Loss of Lgl function was accomplished by either down regulating the *lgl* transcripts through RNAi using the *MS1096* or *Act-GAL4* driver or by inducing FLP-FRT mediated *lgl^−^* mutant somatic clones in *lgl^+^/lgl^−^* background using the MARCM technique (Lee and Luo, 1999; 2001; Gangwani et al., 2020). Since the *lgl^−^* mutant somatic clones by themselves do not grow well due to competitive disadvantage (Agrawal et al., 1995; Menéndez et al., 2010), tumor co-operators like Yorkie or activated Ras were co-overexpressed in the induced *lgl^−^* clones. Following the standardization of the experimental set up, the spatial patterns of expression of the major Hsps, viz., Hsp83, Hsp70, Hsc70, Hsp60, Hsp27 and HSF were examined at different time points after induction of the tumorigenic cells.

Our study showed that cellular levels of the constitutively expressed Hsps like the Hsp83, Hsp60, Hsp27 and Hsc70 were elevated since the early stage of tumorous clones, and these proteins continued to remain high during the entire course of tumor progression. In contrast, the Hsp70, which is an exclusively stress inducible chaperone (Velazquez and Lindquist, 1984; Arya et al., 2007) was expressed only at a later stage. Our model of inducing tumor at a defined time point revealed, for the first time, that levels of the constitutively expressed Hsps are elevated since the early stage of development of a cancer clone. As a proof of principle that the early elevation in levels of these Hsps is necessary for the progression of tumorigenesis, we also show that reduction in levels of Hsp83 from the beginning stage of tumor growth arrested the neoplastic transformation of *lgl^−^* or *lgl^−^* clones co-overexpressing Yorkie (*lgl^−^ yki^OE^*) in wing imaginal discs. Strikingly, HSF, the master regulator of Hsp’s induction during cell stress (Calderwood and Murshid, 2017; Meng et al., 2010; Arya et al., 2007), neither showed major alteration in its levels nor its down-regulation significantly affected the tumorous growth of *lgl^−^ yki^OE^* clones. However, HSF was found to be necessary for the competitive survival of *lgl^−^* clones. Given the conservation of fundamental developmental and disease mechanisms, this model will be very useful for analysing cell processes modulated by the Hsps during different stages of tumorigenesis triggered by various mis-regulated pathways and thus will be of wider relevance in understanding of development of cancer cells and appropriate therapeutic agents.

## Materials and Methods

### Fly stocks and genetics

All fly stocks and crosses were maintained on standard food, containing agar-maize powder, yeast and sugar, at 24±1 °C. The following fly stocks were used to set up crosses to obtain progeny of the desired types: 1. *w*; lgl*^4^ *FRT40A/CyO; Sb/TM6B* (*lgl*^4^ is referred to as *lgl^−^*); 2. *y w* hsFLP; FRT40A w^+^ y^+^/CyO-GFP;* 3. *y w* hsFLP tubGAL4 UAS-GFP; tubGAL80 FRT40A/CyO-GFP; +/+*; 4. *w*; lgl^−^ FRT40A UAS-yki /CyO-GFP;* +/+ (*UAS-yki* as *yki^OE^*); 5. *w*; lgl^−^ FRT40A UAS-Ras^V12^/CyO-GFP* (*UAS-Ras^V12^* referred to as *Ras^v12^*); 6. *MS1096-GAL4; +/+; +/+* (Capdevila and Guerrero, 1994); 7. *y^1^ sc* v^1^ sev^21^; UAS-lgl-RNAi* (BDSC-35773, the *UAS-lgl-RNAi* transgene is referred to as *lgl^RNAi^*); 8. *w*; +/+; UAS-yki^Act^* (BDSC-28817, this transgene is referred to as *yki^Act^*); 9. *y^1^ w^67c23^; UAS-Hsp83* (BDSC-19826, this transgene is referred to as *Hsp83^OE^*); 10. *w^1118^; +/+; Hsp83-RNAi* (NIG 1242R-1, this transgene is referred to as Hsp83^RNAi^); 11. *y^1^ v^1^; UAS-HSF-RNAi/TM3, Sb^1^* (BDSC-27070, this transgene is referred to as *HSF^RNAi^*); 12. *y^1^ v^1^; UAS-HSF-eGFP;* this transgene is referred to as *HSF^OE^*; and 13. *w*; Act5c GAL4/ CyO-GFP; +/+* (Yepiskoposyan et al., 2006)). The stocks 1-6 above were provided by Prof. Pradip Sinha (IIT Kanpur; see (Khan et al., 2013)), 12 was shared by Prof. J. T Lis (USA), while others were either available in our laboratory or were obtained from the Bloomington Drosophila Stock Centre (BDSC) or the National Institute of Genetics (NIG-FLY, Japan).

### Generation of tumors in wing imaginal discs

With a view to generate epithelial tumors in larval wing imaginal discs, *lgl^−^* mutation or *lgl^RNAi^* was used to disrupt cell polarity. In addition, transgenes like *Ras^v12^* or *yki^Act^*, or yki^*OE*^, that express conditionally when driven by an appropriate GAL4 driver, were also used. Two strategies were used for tumor induction. In the first, *MS1096-GAL4* or *Act-GAL4* females were crossed with males of the desired genotype to drive expression of the desired *UAS* carrying transgene in the wing pouch region (Capdevila and Guerrero 1994) or ubiquitously in all cells (Yepiskoposyan et al., 2006), respectively. Synchronized first instar progeny larvae were collected during a 2-hour larval hatching window at 24±1□ and grown at 24±1□ till late third instar stage. Samples of tumorous wing discs were collected by dissecting the desired progeny larvae at different time-points (counted as days After Egg laying or d AEL with 1 day of embryonic development added to the number of days as larvae). In the 2^nd^ approach, we used the MARCM (Mosaic Analysis with a Repressible Cell Marker) technique, which allows Flip-recombinase (FLP) mediated somatic recombination at the Flip-recombinase target (FRT) sites to generate *lgl^−^* homozygous cell clones in *lgl^−^/+* genetic background. Since the somatic recombination event that results in generation of *lgl^−^* homozygous cell also causes loss of the *tubGa180* transgene, the *tubGal4* driver will induce expression of any USA-carrying transgene, including the *UAS-GFP* in the clones, while the other cells *(lgl^−^/+)* remain GFP-negative (Lee and Luo, 1999; 2001).

### MARCM Mosaic analysis

Using the above listed stocks, appropriate crosses were made to finally obtain progenies of the following genotypes.

1. *y w hsFLP tubGAL4 UAS-GFP; tubGAL80 FRT40A/ lgl^−^ FRT40A; +/+* (to generate *lgl^−^* clones in *lgl^+^/lgl^−^* background).
2. *y w hsFLP tubGAL4 UAS-GFP; tubGAL80 FRT40A/ lgl^−^ yki^OE^ FRT40A; +/+* (to generate *lgl^−^ yki^OE^* clones in *lgl^+^/lgl^−^* background).
3. *y w hsFLP tubGAL4 UAS-GFP; tubGAL80 FRT40A/ lgl^−^ Ras^v12^ FRT40A; +/+* (to generate *lgl^−^Ras^V12^* clones in *lgl^+^/ lgl^−^* background).
4. *y w hsFLP tubGAL4 UAS-GFP; tubGAL80 FRT40A/ lgl^−^ FRT40A; Hsp83^RNAi^/+* (to generate *lgl^−^* clones with down-regulated Hsp83 in *lgl^+^/lgl^−^* background).
5. *y w hsFLP tubGAL4 UAS-GFP; tubGAL80 FRT40A/ lgl^−^ yki^OE^ FRT40A; Hsp83^RNAi^/+* (to generate *lgl^−^ yki^OE^* clones with down-regulated Hsp83 in – *lgl^−^ yki^OE^* background).
6. *y w hsFLP tubGAL4 UAS-GFP; tubGAL80 FRT40A/ lgl^−^ FRT40A; Hsp83^OE^/+* (to generate *lgl^−^* clones with up-regulated Hsp83 in *lgl^+^/ lgl^−^* background).
7. *y w hsFLP tubGAL4 UAS-GFP; tubGAL80 FRT40A/ lgl^−^ yki^OE^ FRT40A; HSF^RNA i^ /+* (to generate *lgl^−^ yki^OE^* clones with down-regulated HSF in *lgl^+^/ lgl^−^* background).
8. *y w hsFLP tubGAL4 UAS-GFP; tubGAL80 FRT40A/ lgl^−^ FRT40A; IISF^OE^/+* (to generate *lgl^−^* clones with up-regulated HSF in *lgl^+^/ lgl^−^* background).
9. *y w hsFLP tubGAL4 UAS-GFP; tubGAL80 FRT40A/ lgl^−^ yki^OE^ FRT40A; HSF^OE^ /+* (to generate *lgl^−^ yki^OE^* clones with up-regulated HSF in *lgl^+^/ lgl^−^* background).
10. *y w hsFLP tubGAL4 UAS-GFP; tubGAL80 FRT40A/ w^+^ y^+^ FRT40A; +/+* (to generate *w^+^ y^+^* clones in *y w* background).

The *hsFLP* transgene in the progeny larvae was activated to induce FRT-mediated somatic recombination by heat shocking them at the desired stage of development for a defined time (see Results and Fig. 1) at 37^0^C. In order to maintain uniformity in the heat shock treatment across different samples, 25 progeny first instar larvae were collected within six hours of hatching (a longer window of six hr was taken in this case to obtain a larger number of progeny from which sufficient number of larvae of the desired genotype could be obtained) and transferred to food vials containing *exactly 2ml* of standard food and allowed to grow to the desired age (expressed as hours After Egg Laying (hr AEL), which is the sum of 24 hr of embryonic development and the duration, in hr, of larval development since the collection of 1^st^ instar larvae) when they were heat shocked by adding 0.2ml of standard phosphate buffered saline (PBS), pre-warmed to 37^0^C, into each food vial with larvae (Khan et al., 2013) followed by immediate transfer of the vials to a water bath maintained at 37^0^C for the desired duration (see below). Following heat shock, the food vials with larvae were transferred back to 24^0^C and allowed to grow for defined period of time (see Results). Pupation of the progeny larvae in MARCM crosses was delayed up to 24hr. The larvae carrying the induced clones were identified by screening under a 4x objective on a Nikon E800 fluorescence microscope for absence of GFP expression in the proventriculus region (indicating absence of the *CyO-GFP* balancer), and a variable presence of GFP expressing fluorescent patches; these patches indicated presence of somatic recombination-dependent generation of the GFP expressing *lgl^−^* clones of cells (Lee and Luo 1999; Lee and Luo 2001). Wing and/or eye-antennal imaginal discs were dissected at different hours after the heat shock (hours After Clone Induction or hr ACI) to examine the induced somatic clones.

**Fig. 1.**
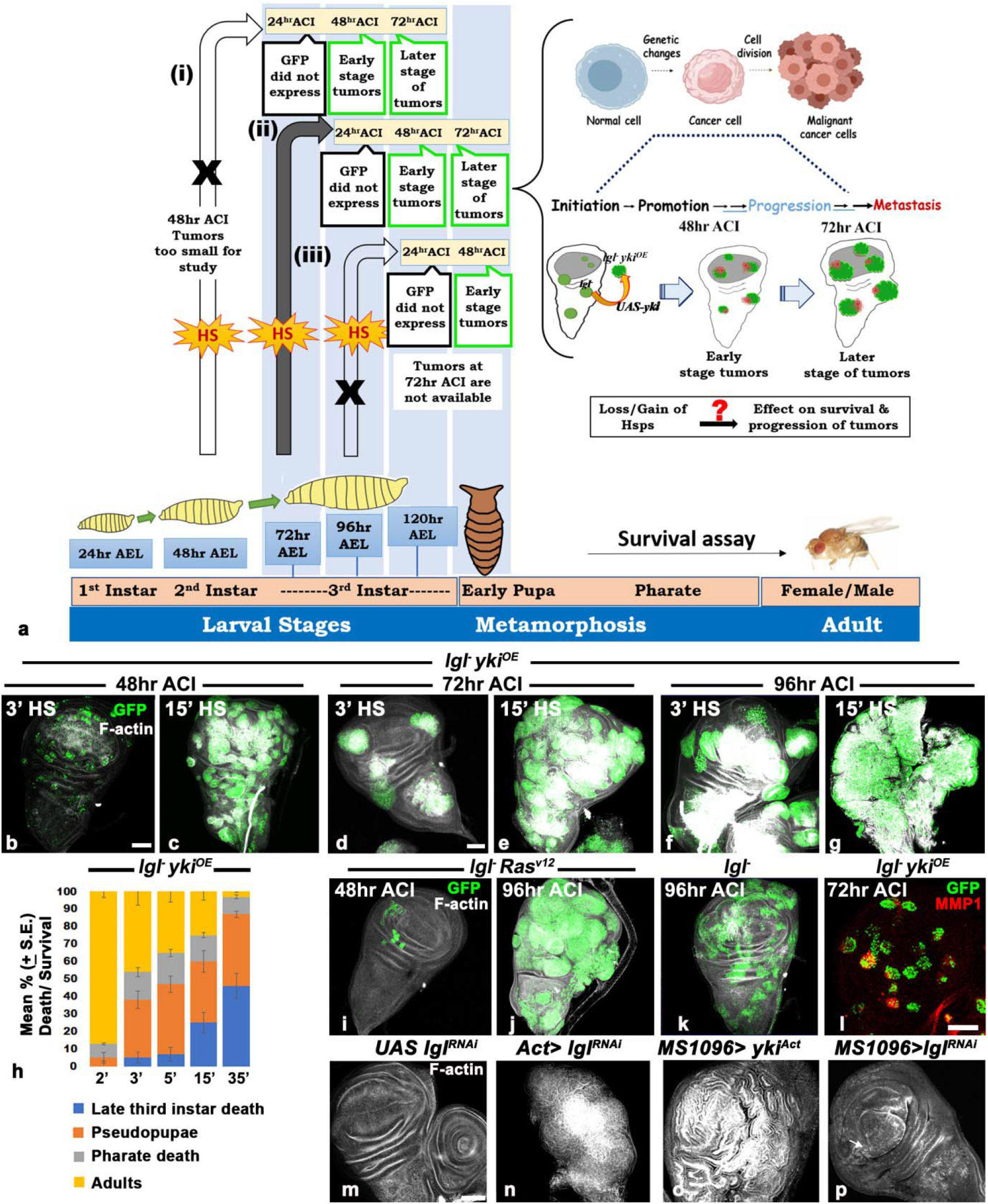
Standardization of developmental stage of larvae and duration of HS for induction of tumors in imaginal discs. **(a)** A schematic of our strategy for generation of clones following HS at different developmental stages and how it affected the analysability of tumor progression and survival of the host. (**b-g**) Confocal projection images of wing discs from larvae exposed to 3min (**d** and **f’**) or 15min (**c, e** and **g’**) HS at 72hr AEL showing combined GFP (green) and F-actin (white) fluorescence in *lgl^−^ yki^OE^* clones at different times ACI (noted on top). **h**, Histograms of mean percentage (+ S.E.) of death/survival till different developmental stages (**Y-axis**) when 72hr old larvae (AEL) were exposed to different durations of HS (in min, **X-axis**) to induce *lgl^−^ yki^OE^* somatic clones. (**i-k**). Confocal projection images of wing discs at different times ACI (noted on left upper corner in each case) following 5min (**i-j**) or 35min (**k**) HS to 72hr old larvae (AEL) showing GFP expressing (green) and F-actin (white) fluorescence in *lgl^−^ Ras^v12^* (**i-j**) or *lgl^−^* (**k**) clones. (**l**) Confocal optical section of wing disc at 72hr ACI showing GFP positive (green) *lgl^−^ yki^OE^* clones induced by 3min HS to 72hr old larva (AEL) some of which are MMP1positive (red). (**m-p**) Confocal projection images of Phalloidin stained (F-Actin, white) wing discs from day 5 (**m**) or day 6 (**n--p**) larvae (AEL) of different genotypes noted above each panel. The scale bars in **b, d** and **l** indicate 50μm; those in **b** and **d** apply to **b** and **c** and to **d-p** (except **l**), respectively.

### Immunostaining

Wing or eye imaginal discs were dissected out in Poels’ salt solution (Tapadia and Lakhotia, 1997) from third instar larvae of the desired genotypes and age (hr ACI), and processed for immunostaining as described earlier (Ray et al., 2019). The following primary antibodies were used: 1. Anti-Hsp27 (Abcam-ab49919; 1:40 dilution) raised in mouse; 2. Anti-Hsp60 (CST-SP60, D307#4870; 1:50 dilution) raised in rabbit; 3. Anti-Hsp70 (7Fb, a kind gift by Prof. M. B. Evgen’ev; 1:200 dilution) raised in rat, which exclusively detects the stress inducible *Drosophila* Hsp70 (Velazquez and Lindquist, 1984); 4. Anti-Hsp/Hsc70 (3.701, a kind gift by Prof. S. Lindquist; 1:50) raised in rat to detect both inducible Hsp70 as well as the constitutive Hsc70 proteins (Velazquez and Lindquist, 1984); 5. Anti-Hsp83 (a kind gift by Prof. Robert Tanguay, 3E6; 1:50 dilution) raised in mouse (Morcillo et al 1993); 6. Anti-cleaved Caspase-3 (Cell Signaling, Asp-216, 1:100 dilution); 7. Anti-MMP1 (DSHB-5H7B11, 1:100); 8. Anti-HSF (a kind gift by Prof. J. T. Lis; 1:300); 9. Anti-Lamin C (DSHB-LC28.26, 1:50). Appropriate secondary antibodies conjugated either with Cy3 (1:200 dilution, Sigma-Aldrich, India) or Alexa Fluor 546 (1:200 dilution; Molecular Probes, USA) were used to detect the given primary antibody. F-Actin was labelled using Phalloidin-633 (#A22284, Invitrogen). 4’, 6-diamidino-2-phenylindole dihydrochloride (DAPI, 1 μg/ml) was used to counterstain DNA. The immunostained tissues were mounted in 1,4-Diazabicyclo [2.2.2] octane (DABCO) antifade mountant for confocal microscopy with Zeiss LSM Meta 510 system using Plan-Apo 20X (0.8 NA), 40X (1.3 NA) or 63X (1.4 NA) oil immersion objectives. Quantitative estimates of proteins in different regions of wing disc or eye discs were obtained using the ZEN (Blue) software. All images were assembled using the Adobe Photoshop 7.0. Significance testing between the data for control and experimental genotypes was performed with the Mann–Whitney U-test or Chisquared test using GraphPad Prism 8.4.3.

### Photomicrography of adult eyes/wings

External morphology of eyes or wings of adult flies of the desired genotype were examined and photographed using a Nikon Digital Sight DS-Fi2 camera mounted on Nikon SMZ800N stereo binocular microscope.

### Survival assay

In order to study the effect of tumorous clones on survival of the host, the larvae were allowed to grow at 24 □ till adults emerged. Numbers of those dying at early pupal (pseudopupae) or late pupal (pharate) stages and adults that emerged were counted. Different data sets were compared with Student’s t-Test for statistical significance using Sigma plot Software.

## Results

### Standardization of heat shock (HS) treatment and the larval developmental stage for induction of tumorous clones in imaginal discs

In pilot experiments to define the timeline when a normal cell begins to develop as tumorous cell, we induced *lgl^−^* or *lgl^−^ yki^OE^* or *lgl^−^ Ras^V12^* clones in *lgl^+^*/*lgl^−^* background by applying HS for 15min at 48hr or 72hr or 96hr AEL (**Fig. 1a**) and examined them at 48hr or 72 or 96hr ACI. Heat shock to 48hr old (AEL) larvae produced many clones which hampered clonal analysis at 72hr or later time ACI. It may be noted that when 72hr old (AEL) larvae were heat shocked and their discs examined at 24hr ACI, the GFP expressing clonal cells in any genotype were nearly completely undetectable (**Fig. 1a**). Since heat shock at the 72hr AEL larval age generated discrete clones and permitted examination of clone-carrying discs at 48hr or 72hr or 96hr ACI, this stage was selected for induction of somatic clones in all the MARCM genotypes (**Fig. 1**). It may be noted that because of the general delay in pupation of the MARCM cross progeny larvae (see Material and Methods), their imaginal discs could also be examined at 96hr ACI (or later if the tumor carrying larvae failed to pupate), when the clones showed massive F-actin accumulation. The 15min HS generated too many and often fused *lgl yki^OE^* or *lgl Ras^V12^* clones, especially at 72hr or 96hr ACI. In parallel, the survival assay also revealed that there was greater pupal and pharate stage death when *lgl^−^ yki^OE^* clones were induced at 72hr AEL by 15 (N=250) or 30min (N=180) HS (**Fig. 1i**). In contrast, a 5min HS provided discrete clones in most discs at 48hr and 72hr ACI, while at 96hr ACI, the discs were nearly completely filled with GFP-expressing tumorous *lgl yki^OE^* or *lgl Ras^V12^* cells (**Fig. 1b-h**). Further experiments showed that HS for 2min or 3min (**Fig. 1b**) to 72hr old larvae induced fewer but discrete *lgl^−^ yki^OE^* clones which could be traced distinctly up to 72 or even 96hr ACI; this also resulted in reduced mortality of host larvae before or after pupation (**Fig. 1i**).

These pilot studies indicated that compared with *lgl^−^ Ras^v12^* clones (**Fig. 1i-j**), the *lgl^−^ yki^OE^* clones showed more aggressive growth during early stages (compare **Fig. 1b** and **1i**). Because of a better consistency in the number and sizes of induced clones, a 3min HS treatment to 72hr old larvae (AEL) was used in subsequent studies to generate *lgl^−^ yki^OE^* clones while a 5min HS treatment at 72hr AEL was used for generating the *lgl^−^ Ras^v12^* clones. As expected, compared to *lgl^−^ yki^OE^* or *lgl^−^Ras^v12^* clones, very few of the *lgl^−^* clones induced by a 5min or 15min HS survived to be visible at 48hr or later ACI time periods as they fail to compete with wild type surrounding cells (Agrawal et al., 1995; Menéndez et al., 2010). A 35min HS at 72hr AEL, however, provided analysable *lgl^−^* clones at 48hr or 72hr or 96hr ACI (**Fig. 1k**). Therefore, the 35min HS was used in subsequent experiments for induction of *lgl^−^* clones.

Since the GFP expressing clones became distinctly visible at 48hr ACI, it was considered as early stage, between initiation and promotion. Many clones showed F-actin accumulation (**Fig. 1d, l**) at 72hr ACI with some of them also showing MMP1 expression (**Fig. 1l**), which indicated that the tumorous clones acquired neoplastic properties by 72hr ACI (Miles et al., 2011).

We also used the *MS1096-GAL4*, which is expresses only in the wing pouch region of wing disc (Capdevila et al., 1994) and the ubiquitously expressing *Act-GAL4* (Yepiskoposyan et al., 2006) drivers to conditionally express *lgl^RNAi^* or *yki^Act^* transgenes and examined the wing discs on day 5 or 6 AEL. *Act-GAL4>lgl^RNAi^* or *MS1096-GAL4>yki^Act^* expression resulted in formation of massively tumorous wing discs (**Fig. 1n-o**), but the *MS1096-GAL4>lgl^RNAi^* expression caused only deformed wing discs without causing a tumorous condition (compare discs in **Fig 1m** and **p**).

These pilot studies also confirmed the varying propensities of progression of the *lgl^−^* clones in different regions of the wing imaginal discs in relation to their genetic background. As reported earlier (Khan et al., 2013), the surviving *lgl^−^* clones were predominantly restricted to the notum area of wing discs. On the other hand, the *lgl^−^ yki^OE^* clones (N= 40 wing discs (WD)) were seen all over the WD but the *lgl^−^ Ras^V12^* clones appeared more often in the wing pouch region since in 28% of 32 WD clones were seen only in the pouch region, while in the remaining discs, a few clones were also present in the notum and hinge regions. Interestingly, despite the initially slower growth of *lgl^−^ Ras^V12^* clones in comparison to *lgl^−^ yki^OE^*, the larvae carrying *lgl^−^ Ras^V12^* clones (N=66) showed greater lethality than seen in *lgl^−^ yki^OE^* clone-carrying larvae (N=78).

### Tumorous cells over-express Hsp83 at the early stage but non-tumorous neighbouring cells also exhibit elevated Hsp83 at later stages of tumor growth

We examined spatial and temporal expression of Hsp83, the *Drosophila* homologue of mammalian Hsp90, in different tumor backgrounds (**Fig. 2**). Enhanced levels of Hsp83 in the *lgl^−^ yki^OE^* neoplastic clonal cells were observed as early as 48hr ACI (**Fig. 2a-a’**), with markedly enhanced levels in cells at the periphery of each clone, creating a ring like appearance (**Fig. 2e-e’**). Hsp83 was mostly cytoplasmic and became uniformly high all through the clonal area at later stages (**Fig. 2b-c’**). Interestingly, the Hsp83 levels were also enhanced in the neighbouring non-tumorous cells, especially at 72hr ACI and later, when the F-actin accumulation in the clones became increasingly high (**Fig. 2b’, e-f’**). The *lgl^−^ Ras^v12^* neoplastic clones also exhibited a similar pattern of Hsp83 expression during their progression (expression at 96hr ACI is shown in **Fig. 2d-d’**). The Hsp83 levels in wing discs from *Act>lglRNAi* or *MS1096>yki^act^* 5- or 6-day old larvae, respectively, were also enhanced with high F-actin accumulation all through the disc (**Fig. 2e-f’**).

**Fig 2.**
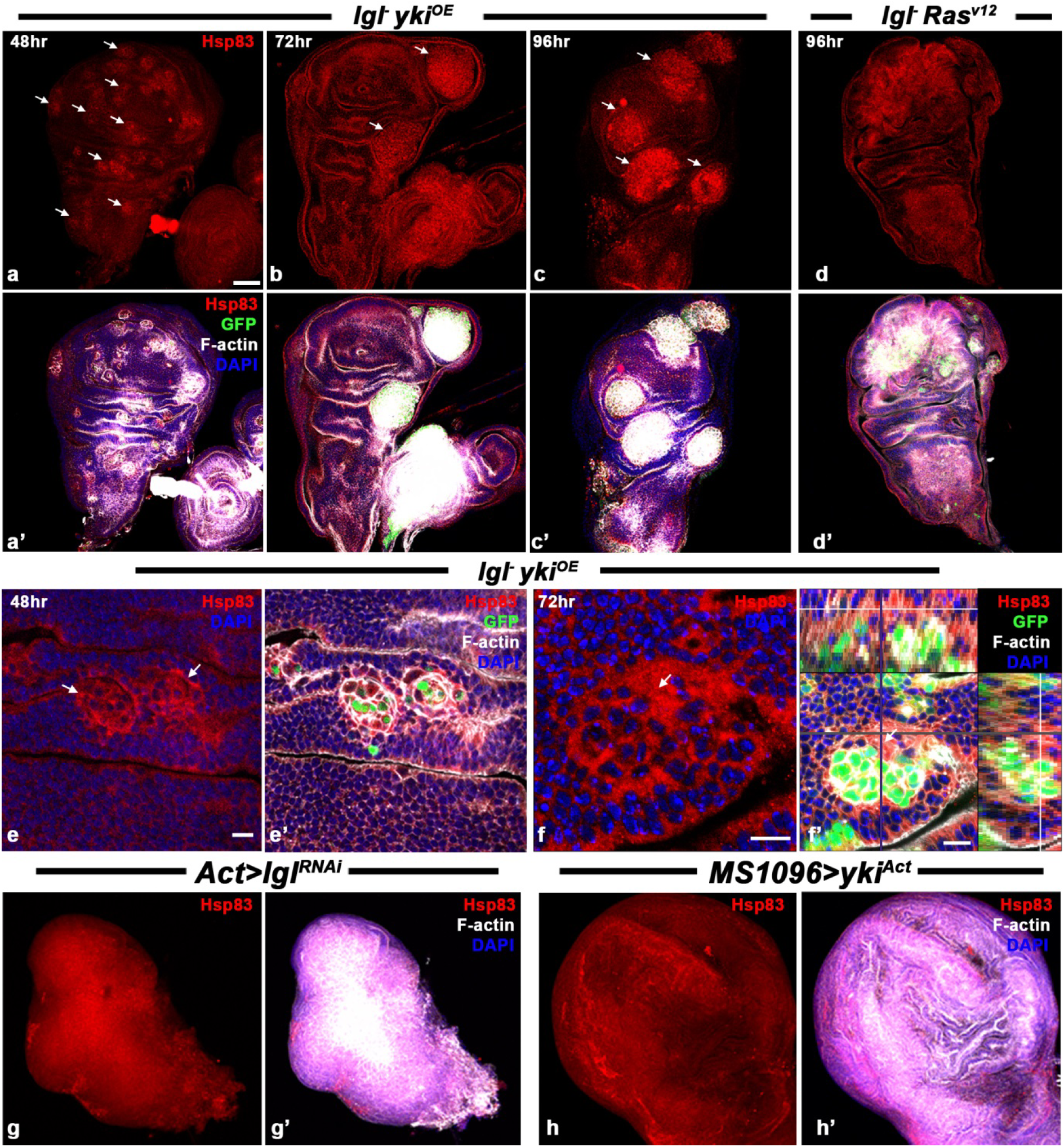
Hsp83 is elevated in tumorous and non-tumrous neighboring cells as the tumor progresses. **(a-d’)** Confocal optical sections of wing discs showing Hsp83 (red) over-expression in *lgl^−^ yki^OE^* **(a-c’)** or *lgl^−^ Ras^v12^* (**d-d’**) clones (white arrows) induced by 3min (**a-c’**) or 5 min (**d-d’**) HS to 72hr old (AEL) larvae at 48hr (**a-a’**), 72hr (**b-b’**) and 96hr (**c-d’**) ACI. **a’-d’** show combined Hsp83 (red), GFP (green), F-actin (white) and DAPI (blue) fluorescence in discs shown in **a-d**, respectively. (**e, f**) higher magnification images to show elevated Hsp83 (red) levels in non-GFP expressing cells (white arrows in **e** and **f**) adjacent to *lgl^−^ yki^OE^* clone (green); **f’** is an orthogonal image of the disc in **f** with Y-Z and X-Z projections shown on top and right, respectively; **e’** and **f’** show combined Hsp83 (red), GFP (green), F-actin (white) and DAPI (blue) fluorescence; white arrows in **f-f’** indicate a non-GFP expressing cell with elevated Hsp83. (**g, h**) Confocal projection images of *Act-GAL4>lglRNAi* (**g-g’**) or *MS1096-GAL4>yki^Act^* (**h-h’**) tumorous wing discs from day 5 (**g-g’**) or day 6 (**h-h’**) larvae (AEL) showing elevated Hsp83 throughout the disc (red); **g’** and **h’** show combined Hsp83 (red), F-actin (white) and DAPI (blue) fluorescence. Scale bar in **a** denotes 50μm and applies to **a-d’** and **g-h’**, the scale bars in e-f’ denote 10μm.

### Tumor cells over-express HSC70 from the early stage but the stress-inducible Hsp70 is expressed only in a few cells of the later stage tumorous *lgl^−^ yki^OE^* clones

Proteins belonging to the Hsp70 family have both contitutive as well as inducible forms and all of them have been implicated in tumorigenesis in mammals (Zaarur et al., 2006; Meng et al., 2010). In *Drosophila*, the inducible form is the Hsp70 while the several constitutive forms are grouped as Hsc70 (Velazquez and Lindquist, 1984; Elefant and Palter, 1999). We looked at expression of both classes of this family of proteins in clones, using two different antibodies. The 7Fb antibody detects only the stress inducible *Drosophila* Hsp70 while the 3.701 antibody detects both inducible as well as constitutive forms, i.e., Hsp70 as well as the various Hsc70 proteins (Velazquez and Lindquist, 1984). The difference between the staining with 3.701 and 7Fb can thus be ascribed to Hsc70 expression. Staining with 3.701 antibody at 48hr ACI showed slightly increased levels of Hsc70/Hsp70 proteins in *lgl^−^ yki^OE^* clones compared to their neighbouring areas (**Fig. 3a-a’)**. Surprisingly, however, the 7FB antibody did not show any Hsp70 staining in or outside the *lgl^−^ yki^OE^* clones at 48hr ACI (**Fig. 3b-b’**), implying thereby that the increased staining observed with the 3.701 antibody at 48hr ACI reflects enhanced levels of some or all of the various Hsc70 proteins since the early stage of clone growth. The 3.701 antibody showed further increase in abundance across the clone areas at 72hr ACI, more so at the clone periphery (**Fig. 3c-c’**). Interestingly, however, the 7FB antibody staining pattern at 72hr ACI was very different from that of the 3.701 antibody. In a few clones (38 of 106 examined), the entire clone area showed Hsp70 at a low level with some cells showing a much higher Hsp70 staining (**Fig. 3d-d’**, marked with yellow arrows). But in the remaining 68 clones at 72hr ACI, only a few of the *lgl^−^ yki^OE^* clone cells were distinctly 7FB positive (**Fig. 3e-e’**, marked with white arrows) while staining in rest of the clone area was negligible.

**Fig. 3.**
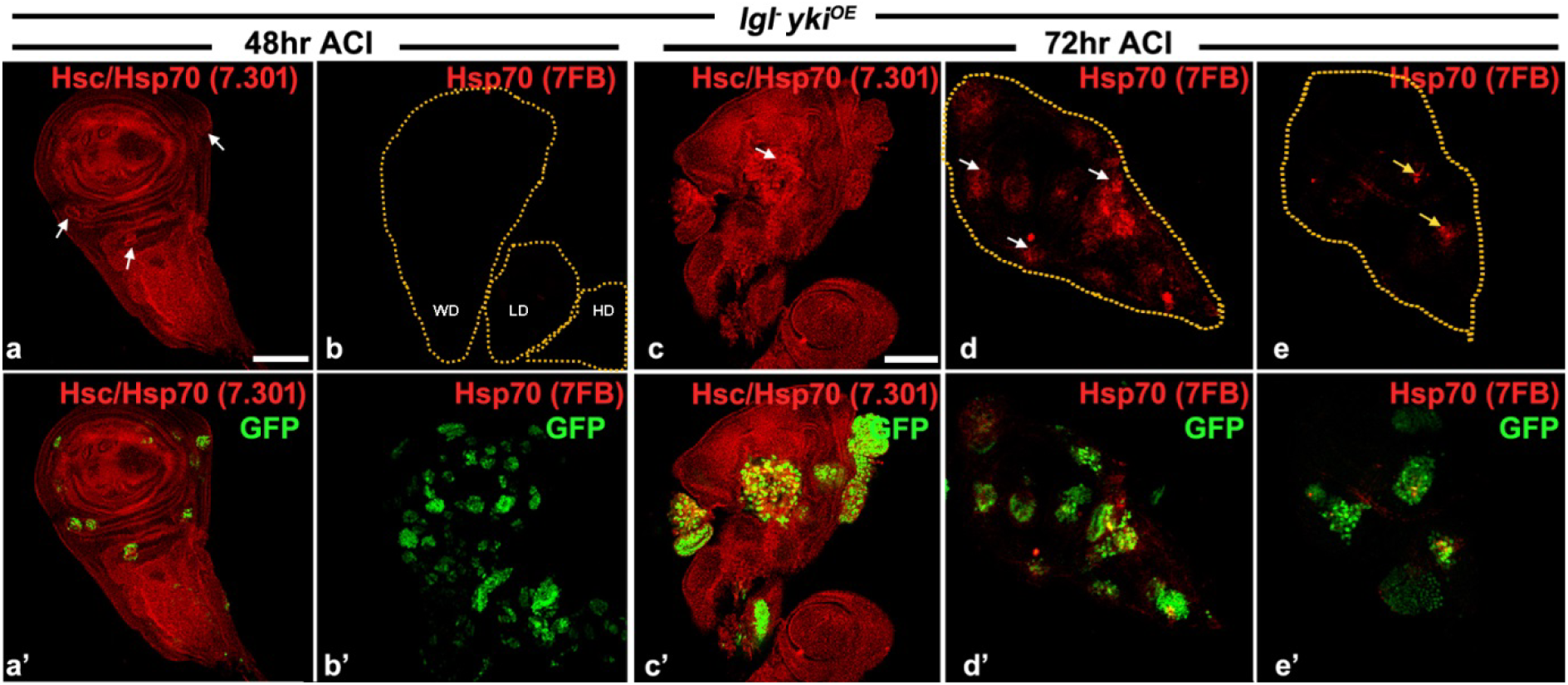
The *lgl^−^ yki^OE^* clonal cells over express Hsc70 since the early stages but Hsp70 is expressed only in a few *lgl^−^ yki^OE^* cells at later stages. (**a-a’** and **c-c’**) Confocal images of wing imaginal discs showing over-expression of Hsc70/Hsp70 (red; white arrows) at 48hr (**a-a’**) and 72hr (**c-c’**) ACI in GFP positive *lgl^−^ yki^OE^* clones (green) induced by 3min HS at 72hr AEL. (**b-b’** and **d-e’**). Wing imaginal discs immunostained with the Hsp70-specific7FB antibody (red) at 48hr (**b-b’**) and 72hr (**d-e’**) ACI; white arrows in **d** and **e** indicate the GFP positive (green) clones seen in **d’** and **e’**, respectively; dotted lines in **b** outline the wing (**WD**), leg (**LD**) and haltere (**HD**) discs; **a’-e’** show combined red (Hsc70/Hsp70) and green (GFP) fluorescence in the discs shown in **a-e**, respectively. The confocal images in **a-a’** and **c-c’** are optical sections while those in **b-b’, d-e’** are projections. Scale bars in **a** and **c** denote 50μm and apply to **a-b** and **c-e’,** respectively.

### Tumorous clones over-express Hsp60 at early stage and also in non-tumorous neighbouring cells at later stages of tumor growth

We examined expression of Hsp60 in different tumor backgrounds (**Fig. 4**). Enhanced levels of Hsp60 in the *lgl^−^ yki^OE^* clonal cells were observed as early as 48hr ACI (**Fig. 4a-a’**), with markedly greater enhancement in cells at the periphery of each clone (**Fig. 4e-e’)**, as also noted above for Hsp83. At later stages, again like the Hsp83, the Hsp60 distribution was uniformly high all through the clonal area (**Fig. 4b-c’**) with high levels in the neighbouring non-tumorous cells as well (**Fig. 4f-f’)**. The *lgl^−^ Ras^v12^* neoplastic clones also exhibited a similar pattern of Hsp60 expression during their progression (expression at 96hr ACI is shown in **Fig. 4d-d’**). The levels of Hsp60 appeared to correlate with the extent of F-actin accumulation in clones at later hr ACI (**Fig. 4**).

**Fig 4.**
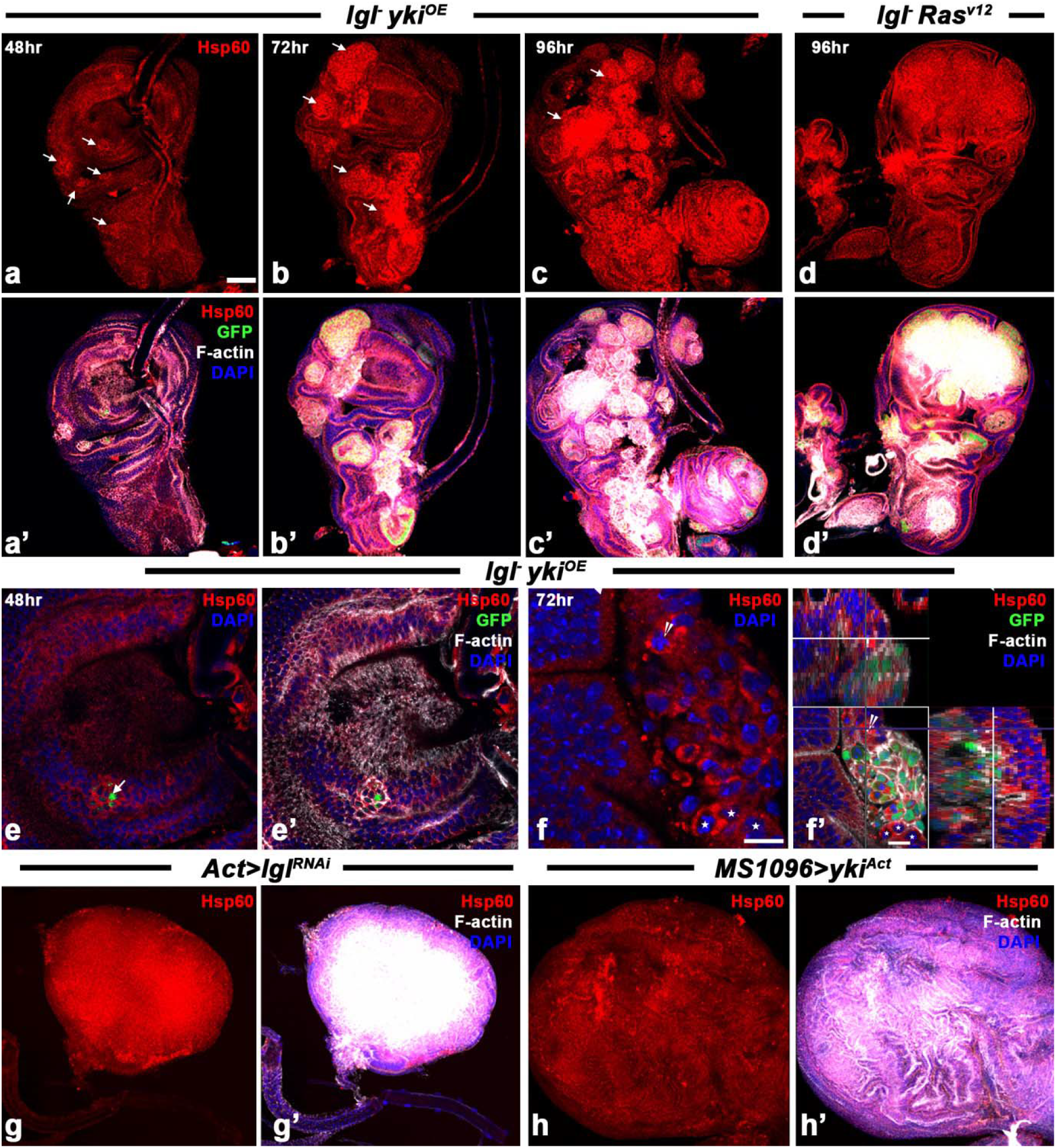
Hsp60 is over-expressed by tumor cells since the beginning of their development till late stages. **(a-c’)** Confocal optical sections showing Hsp60 (red) over-expression in *lgl^−^ yki^OE^* GFP (green) positive clones at 48hr (**a-a’**), 72hr (**b-b’**) and 96hr (**c-c’**) ACI. (**d-d’**) *lgl^−^ Ras^v12^* tumors showing enhanced Hsp60 (red) at 96hr ACI. (**e-f’**) Confocal optical sections of *lgl^−^ yki^OE^* clones at higher magnification to show cytoplasmic accumulation of Hsp60 (red); **f’** is orthogonal image of the region of the wing disc presented in **f** to show elevated Hsp60 in cytoplasm in non-GFP positive cells (white stars in **f-f’**) neighbouring a *lgl^−^ yki^OE^* clone; arrowhead in **f-f’** indicates a GFP-ve cell at the intersection of X-Y axis for the ortho image. **a-f** show only Hsp60 (red) while **a’**-**f’** show combined Hsp60 (red), GFP (green), F-actin (white) and DAPI (blue) fluorescence images of **a-f**, respectively. White arrows in **a-e** indicate clonal areas. (**g-h’**). Confocal projection images of *Act-GAL4>lglRNAi* (**g-g’**) or *MS1096-GAL4>yki^Act^* (**h-h’**) tumorous wing discs showing elevated Hsp60 (red); **g’**. and **h’** show F-actin (white) and DAPI (blue) counterstaining. Scale bar in **a** denotes 50μm and applies to **a-h’**; scale bar in **f** denotes 10μm and applies to **e-f**; scale bar in **f** represents 10 μm.

The tumorous wing discs from *Act>lglRNAi* or *MS1096>yki^Act^* larvae also showed enhanced levels of Hsp60 and Factin all through the disc (**Fig. 4e-f**’).

### Hsp27, a member of the sHsp family, is also over-expressed by tumor cells from the beginning till late stages of their growth

Enhanced levels of cytoplasmic Hsp27 were seen in *lgl^−^ yki^OE^* neoplastic clonal cells at 48hr ACI (Fig. 5a-a’). The Hsp27 levels further increased at later stages, becoming more uniform all through the clone areas (**Fig. 5b-c’**). The *lgl^−^ Ras^V12^* neoplastic cells exhibited a similar pattern of Hsp27 expression during their progression (**Fig. 5d-d’**).

**Fig. 5.**
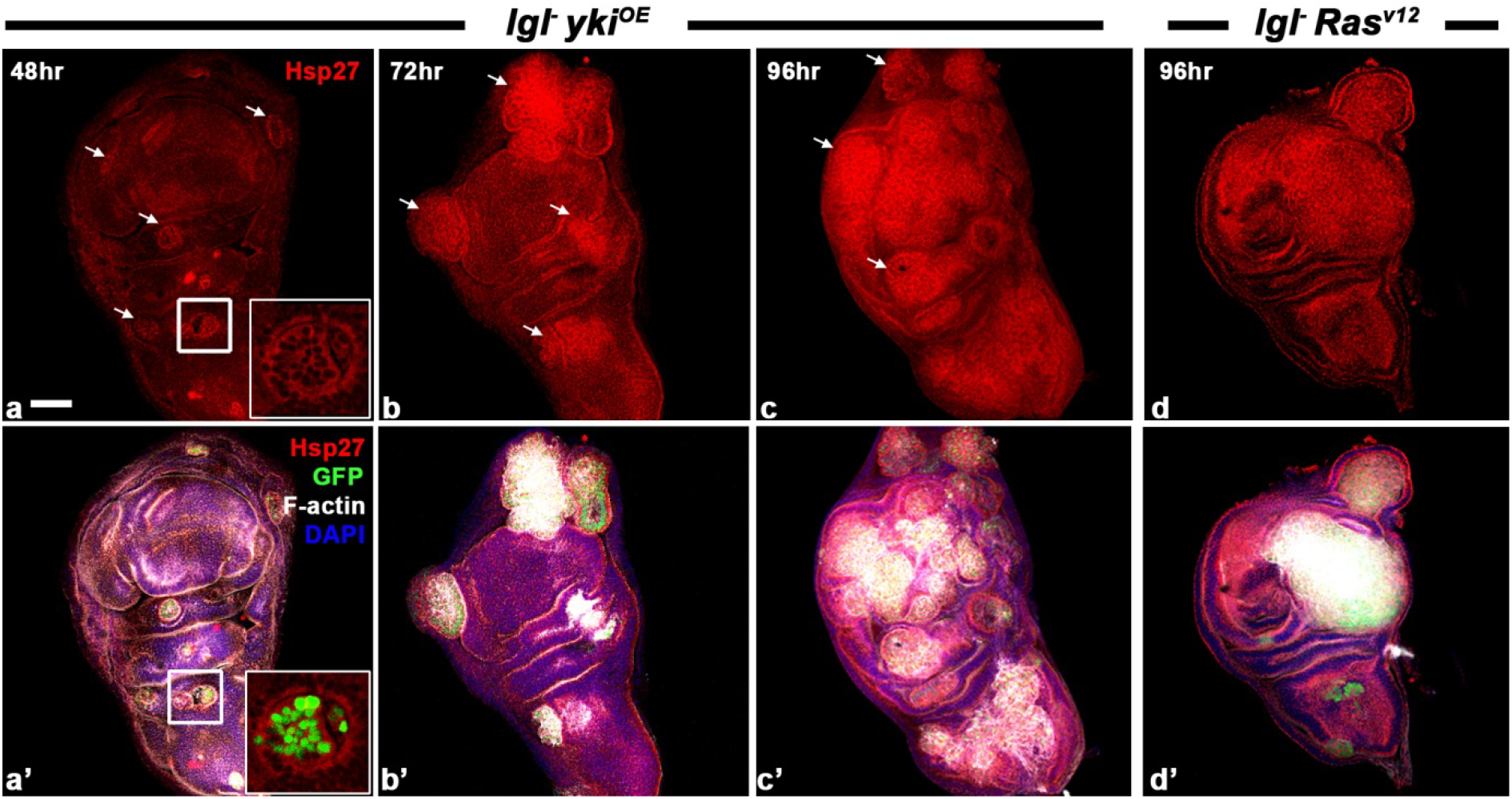
Neoplastic cells over express Hsp27 since the beginning of their development till late stages of progression. (**a-d’**) Confocal optical sections showing over-expression of Hsp27 (red) in *lgl^−^ yki^OE^* (**a-c’**) and *lgl^−^ Ras^V12^* (**d-d’**) clones in wing imaginal discs at diferent time points (noted in left upper corner of each column) ACI. Panels in **a-d** show Hsp27 (red), while those in **a’-d’** show combined Hsp27 (red), GFP (green), DAPI (blue) and Phalloidin (white) fluorescence. White arrows in **a-d** mark tumrous clonal areas. Insets in **a-a’** show enlarged images of boxed areas in **a-a’**, respectively. Scale bar in **a** represents 50μm and applies to all.

### The heat shock treatment used to induce the clones by itself does not elevate Hsp levels

Since we found levels of different Hsps in the clones to be elevated at different time points after heat shock, we also checked if the HS treatment applied to younger larvae could have led to the elevated levels of the Hsps at later time points. Therefore, as a control we examined the expression of Hsp83 and Hsp60 in wing discs of *lgl^−^/lgl^+^* larvae *(y w hsFLP tubGAL4 UAS-GFP; lgl^−^ FRT40A/CyO-GFP; +/+)* exposed to 35 min HS. These discs showed normal development with ubiquitous GFP (because of the *tub-GFP* transgene) expression. All the imaginal discs examined showed Hsp83 (N=12) or Hsp60 (N=16) to be present at the normal low levels (**Fig. 6**). Thus, the above noted elevated levels of Hsps seen in the tumors at 48hr or later ACI are not because of persistence of any Hsps induced by the brief HS applied to larvae for induction of the clones.

**Fig. 6.**
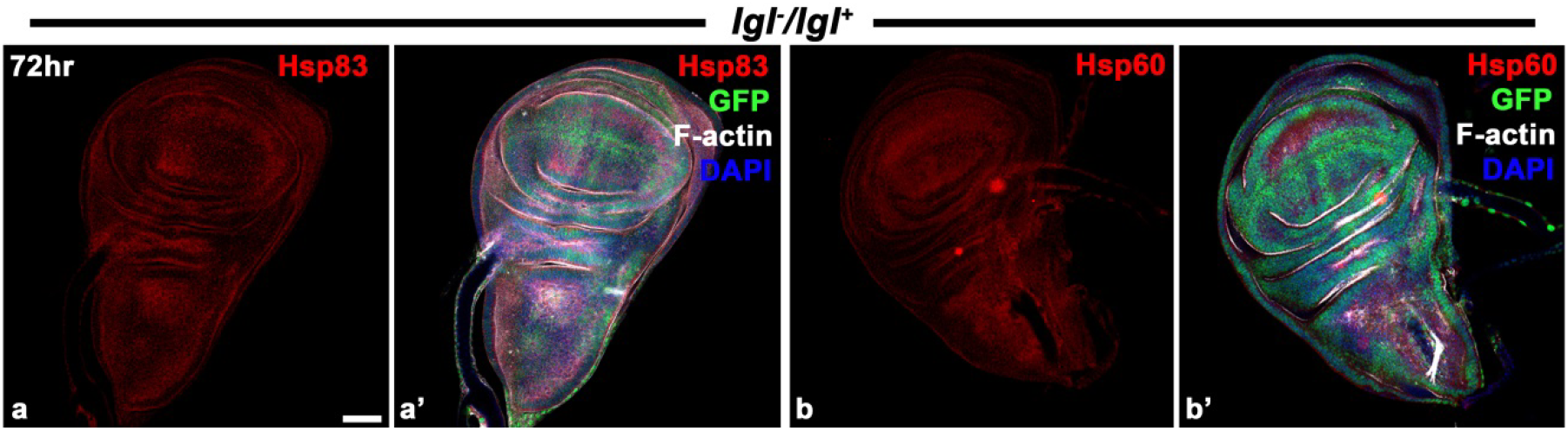
Heat shock to induce MARCM clones does not lead to persistence of any induced Hsp at later time points. (**a-b’**) Wing imaginal discs immunostained for Hsp83 (red, **a-a’**) or Hsp60 (red, **b-b’**) at 72hr after 35min heat shock to 72hr old (AEL) *tub-GFP; lgl^−^/lgl^+^* larvae. Combined Hsp (red), GFP (green), F-Actin (white) and DAPI (blue) fluorescence is shown in **a’** and **b’**.

### Hsp83 is essential for post-initiation survival and progression of tumor cells

To understand the functional relevance of the elevated levels of Hsps in the tumorous clones, we examined growth of *lgl^−^ yki^OE^* clones when Hsp83 levels were down-regulated by co-expressing *UAS-Hsp83^RNAi^* transgene in *lgl^−^ yki^OE^* clones. Interestingly, the *lgl^−^ yki^OE^ Hsp83^RNAi^* clones (N=42 wing discs) showed poor survival at 48hr ACI (**Fig. 7a-b’**), with a significant reduction in their mean frequency when compared with that of *lgl^−^ yki^OE^* clones (N= 57 wing discs; p<0.03 on Mann–Whitney test, **Fig. 7e**). In parallel with the decrease in the frequency of clones/disc in the pouch area, the % Clonal area/disc (measured as the cumulative area covered by all clones in the pouch region of a disc as percent of the total area of the same wing pouch in maximum intensity projection images) of the surviving *lgl^−^ yki^OE^ Hsp83^RNAi^* clones (N= 42 discs) was also significantly smaller than in the parallel *lgl^−^ yki^OE^* controls (N= 57 discs; p<0.05 on Mann–Whitney test, **Fig. 7 f**). At 72hr ACI also, *lgl^−^ yki^OE^ Hsp83^RNAi^* clones in wing discs (N= 41 discs) showed further reduction in % Clonal area/disc when compared with the same age *lgl^−^ yki^OE^* clones (N= 54 discs, p<0. 01 on Mann-Whitney test, **Fig. 7 c-d’, g**). The accumulation of F-actin in *lgl^−^ yki^OE^ Hsp83^RNAi^* clones at 72hr ACI also appeared to be less evident (**Fig. 7d**) than in the same age *lgl^−^ yki^OE^* (**Fig. 7c**) clones. Immunostaining for assessing expression of the DCP-1 protein revealed that a significantly high proportion of the surviving *lgl^−^ yki^OE^ Hsp83^RNAi^* clones (N=41 discs) were DCP-1 positive than in discs with *lgl^−^ yki^OE^* clones (N= 54 discs; **Fig. 7 h**, P<0.01 on Chi-squared test). In agreement with the poor survival of *lgl^−^ yki^OE^ Hsp83^RNAi^* clones, the survival of the host larvae was also better than that of the larvae carrying *lgl^−^ yki^OE^* clones since in the former case, fewer pseudopupae or dying pharates were seen (**Fig. 7i**). Interestingly, an over-expression of Hsp83 in *lgl^−^ yki^OE^ Hsp83^OE^* clones did not have any significant effect since their growth and abundance appeared similar to those of *lgl^−^ yki^OE^* clones at 48hr and later time points after ACI (data not presented).

**Fig. 7.**
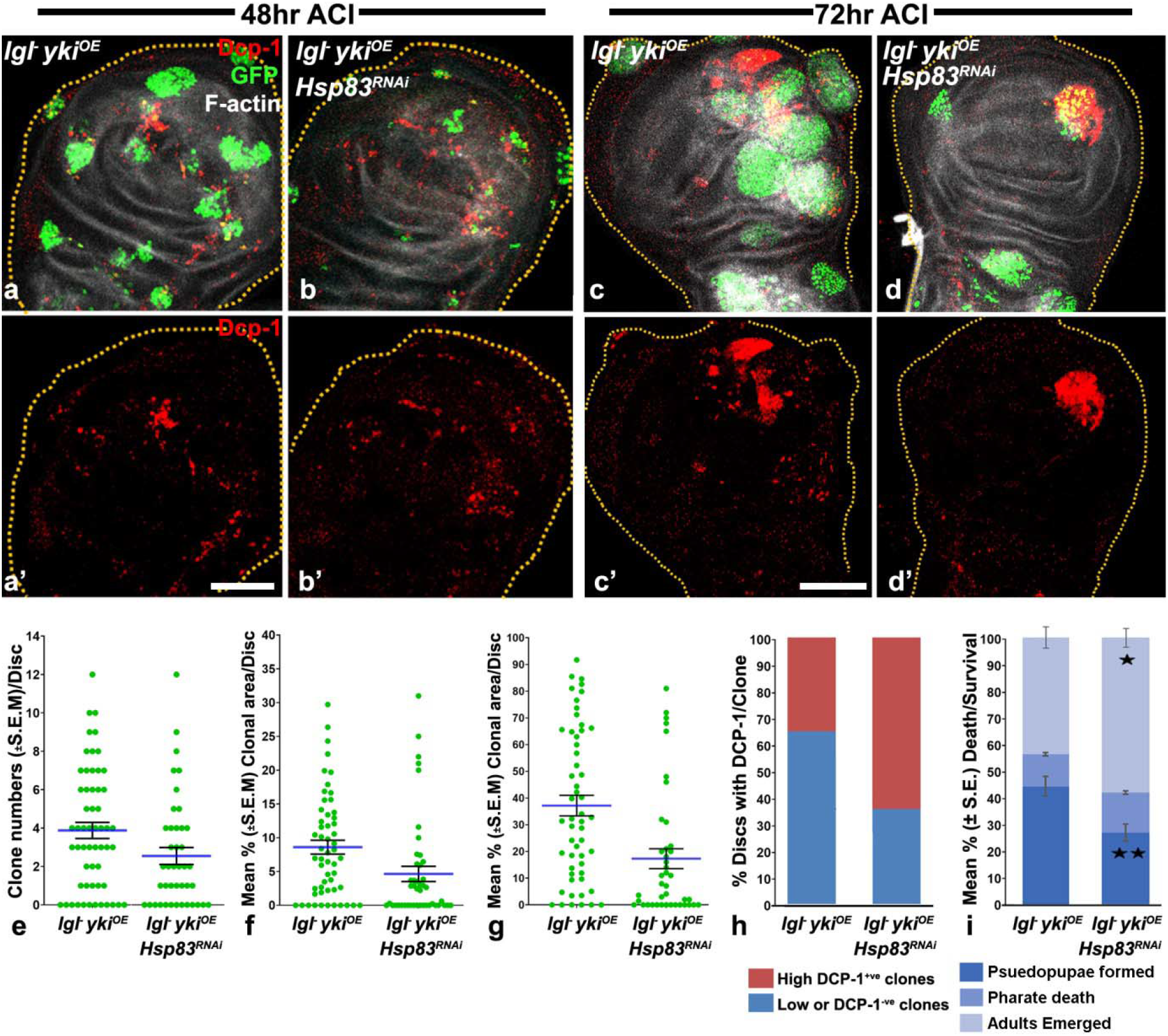
Loss of Hsp83 results in increased death and elimination of *lgl^−^ yki^OE^* clones and better survival of the host. (**a-d’**) Confocal projection images of wing discs (outlines marked with yellow dotted lines) carrying *lgl^−^ yki^OE^* (**a, a’, c, c’**) or *lgl^−^ yki^OE^ Hsp83^RNAi^* (**b, b’, d, d’**) GFP expressing (green) clones showing distribution of Dcp-1 (red) and F-actin (white) at 48hr (**a-b’**) and 72hr (**c-d’**) ACI; scale bar in **a’** represents 50μm and applies to **a-d’**. (**e-g)** are scatter plot graphs with **e** showing mean numbers (±S.E.M.) of clones per disc (Y-axis) at 48hr ACI in different genotypes (X-axis), **f** and **g** showing mean (±S.E.M.) % Clonal Area/Disc (Y-axis) at 48hr (**f**) and at 72hr ACI (**g**) in different genotypes (X axis). (**h**) Bar graph histogram depicting % of wing discs (Y-axis) of different genotypes (X-axis) expressing negligible DCP-1 (blue shaded region, DCP-1-ve) or high (red shaded region, DCP-1 +ve) in GFP positive clones at 72hr ACI. (**i**) Bar histograms show mean (±S.E.) % death/survival at different developmental stages (Y-axis) of the 72hr old larvae exposed to 3’ HS to induce *lgl^−^ yki^OE^* or *lgl^−^ yki^OE^ Hsp83^RNAi^* somatic clones (X-axis); * indicates P<0.05 and ** indicate P<0.01.

Cell-competition regulates the transformation of a normal cell into a neoplastic cell (Baker, 2020). Therefore, we also examined role of Hsp83 in survival and growth of *lgl^−^* clones (generated by a 35min HS at 72hr AEL) in wing and eye discs by co-expressing either *Hsp83^RNAi^* or *Hsp83^OE^* transgene. The *lgl^−^ Hsp83^RNAi^* clones were less frequent than the *lgl^−^* clones at 48hr (data not shown) and at 72hr ACI in wing as well as eye discs (**Fig. 8**). Interestingly, while the abundance and sizes of *lgl^−^ Hsp83^OE^* clones in wing discs at 72hr ACI appeared similar to those of the *lgl^−^* clones (**Fig. 8a, c**), the *lgl^−^ Hsp83^OE^* clones in eye discs appeared larger (**Fig. 8d, f**). In agreement, the survival of larvae carrying *lgl^−^ Hsp83^RNAi^* clones to adult stage was significantly better than those carrying *lgl^−^* clones while that of larvae carrying *lgl^−^ Hsp83^OE^* clones remained more or less unchanged (**Fig. 8g**). A proportion of the flies emerging from larvae carrying *lgl^−^* clones showed wing abnormalities (like blisters, missing vein and/or wing shape changes) and necrotic patches in and around eyes (**Fig. 8i**). In agreement with fewer *lgl^−^ Hsp83^RNAi^* clones and better survival of these larvae to adult stage, the frequencies of wing (**Fig. 8j**) and eye (**Fig. 8k**; N= 85 flies) phenotypes in the surviving flies were also significantly lower than in flies developing from larvae with *lgl^−^* clones (N= 68 flies). On the other hand, frequency of necrotic patches in and around eyes of adults developing from larvae carrying *lgl^−^ Hsp83^OE^* clones (N= 71 flies) was significantly greater than in the adults developing from larvae with *lgl^−^* clones (**Fig. 8k**) although the wing abnormalities in these two groups of flies remained similar (**Fig. 8j**).

**Fig. 8.**
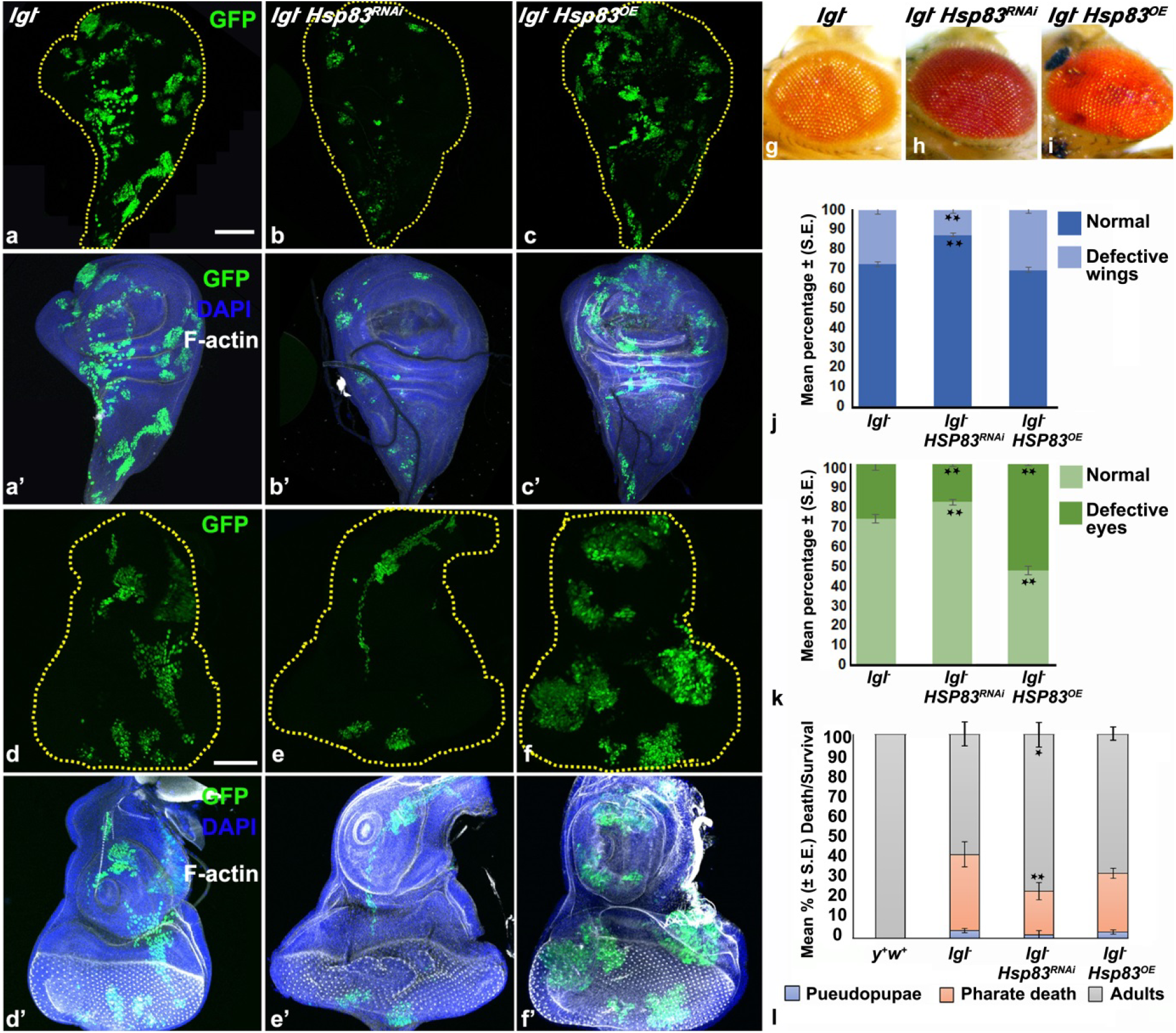
The *lgl^−^* clones require Hsp83 for survival. (**a-f’**) Confocal projection images of wing (**a-c’**) and eye-antennal (**d-f’**) imaginal discs (outlines in **a-f** marked with dotted lines) showing GFP expressing (green) tumorous clones of different genotypes (noted above each column) at 72hr ACI by 35min HS at 72hr AEL; images in **a’-f’** show combined GFP (green), DAPI (blue) and F-actin (white) fluorescence for discs in **a-f**, respectively; scale bars in **a** and **d** represent 50μm and apply to **a-c’** and **d-f’**, respectively. (**g-i**) Photomicrographs of eyes of adults emerging from larvae carrying *lgl^−^* (**g**) or *lgl^−^*; *Hsp83^RNAi^* (**h**) or *lgl^−^; Hsp83^OE^* clones (**i**). (**j**, **k**) Bar histograms of mean percent (+ S.E.) defects observed in wings (**j**) and eyes (**k**) in adults developing from larvae carrying clones of different genotypes (noted below each bar); ** in **j** and **k** indicate P<0.01 when compared with corresponding values for larvae with *lgl^−^* clones. (**l**) Bar histograms show Mean % (+S.E.) survival till different developmental stages (Y-axis) of the 72hr old larvae exposed to 35’ HS to induce *lgl^−^, lgl^−^ Hsp83^RNAi^* or *lgl^−^ Hsp83^OE^* somatic clones (X-axis). The * in (**j-l**) indicate P<0.05 while ** indicate P<0.01 when compared with corresponding values for larvae with *lgl^−^* clones.

### Altered levels of HSF do not affect *lgl^−^ yki^OE^* clone growth but HSF over-expression improves survival and transformation of *lgl^−^* clones

HSF is known to be master regulator of the stress-induced heat shock gene expression and to be also associated with several genes other than Hsps during the development so that it has a significant role in *Drosophila* development (Birch-Machin et al., 2005). HSF has also been reported to play important role in transformation of a normal cell to cancer cell and subsequent tumorigenesis in mammalian cancers (Alasady and Mendillo, 2020; Dai et al., 2007; Zaarur et al., 2006; Meng et al., 2010). Therefore, we also checked effects of down- or up-regulation of HSF in *lgl^−^* and *lgl^−^ yki^OE^* clones on their growth.

Over-expression of HSF in *lgl^−^; HSF^OE^* clones resulted in morphological change of the wing disc due to significant increased survival of *lgl^−^* clones (**Fig. 9a-b**). Compared to the *lgl^−^* clones, elevated levels of HSF helped the *lgl^−^; HSF^OE^* clones in escaping the competitive elimination and to acquire neoplastic properties as observed by F-actin aggregation (N=22) and expression of MMP1 in these clones (N=16)(compare **Fig. 9c-c’** with **d-d’**).

**Fig. 9.**
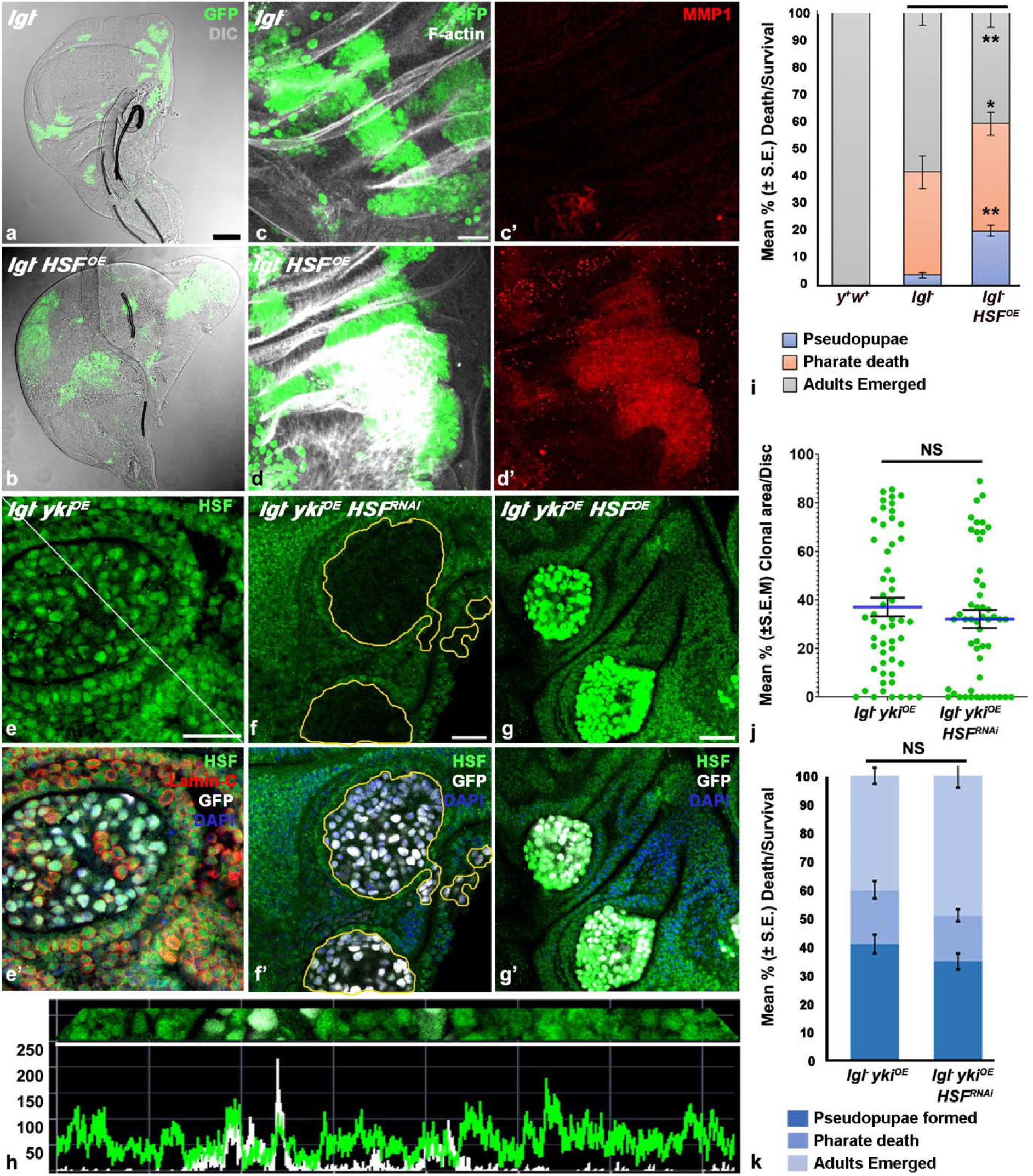
HSF overexpression improves survival of *lgl^−^* clones but HSF down- or up-regulation has little effect on growth of *lgl^−^ yki^OE^* clones. **(a-b)** Confocal projection images of wing imaginal discs showing distribution of GFP positive (green) and morphology of wing discs (DIC) carrying *lgl^−^* (**a**) or *lgl^−^ HSF^OE^* (**b**) clones on day 5 AEL. (**c-d’**) Confocal projection images showing distribution of F-Actin (white, **c** and **d**) and MMP1 (red, **c’** and **d’**) in GFP positive (green) *lgl^−^* (**c-c’**) or *lgl^−^ HSF^OE^* (**d-d’**) clones at 96hr ACI. (**e-g’**) Confocal optical sections of wing imaginal discs showing distribution of immunostained HSF (green), Lamin-C (red, **e’**) in GFP (white) positive *lgl^−^ yki^OE^* clones; (**e-e’**) or *lgl^−^ yki^OE^ HSF^RNAi^* (**f-f’**) or *lgl^−^ yki^OE^ HSF^OE^* (**g-g’**), with clonal areas outlined with dotted yellow lines, showing distribution of HSF (immunostained, green) and GFP (white); **e’-g’** also show DAPI (blue) fluorescence. (**h**) Intensity profile of GFP (white) and HSF (green; Y-axis) fluorescence along the white diagonal line in **e**. dotted lines (X-axis). Note that the GFP signals in **b, d** and **g’** reflect combined GFP fluorescence output of the *UAS-HSF^OE-eGFP^* and *UAS-GFP* transgenes. **(i)** Bar histograms show Mean % (+S.E.) death/survival till different developmental stages (Y-axis) of the 72hr old larvae exposed to 35’ HS to induce *lgl^−^* or *lgl^−^ HSF^OE^* somatic clones (X-axis; *y^+^ w^+^* larvae were used as control). **(j)** Scatter plot graphs showing mean (+S.E.M.) % Clonal Area/Disc (Y-axis) at 72hr ACI in different genotypes (X axis). (**k**) Bar histograms show Mean % (+S.E.) death/survival till different developmental stages (Y-axis) of the 72hr old larvae exposed to 3’ HS to induce *lgl^−^ yki^OE^* or *lgl^−^ yki^OE^ HSF^RNAi^* somatic clones (X-axis). **\*** and **\*\*** in **i** indicate P<0.05 and P<0.01, respectively, between the compared genotypes indicated by horizontal lines on top of the bars; **NS** in **j** and **k** indicate P>0.05 between the compared genotypes indicated by horizontal line on top of the data sets. Scale bar in **a’** represents 50μm and applies to **a-b,** while scale bars in **c, e, f** represent 20μm and apply to **c-d’, e-e’** and **f-g’** respectively.

Since the *lgl^−^ yki^OE^* clones were found to over-express different Hsps from the early stage of tumor growth, we examined HSF levels in these clones and effects of down-or up-regulation of HSF on growth of *lgl^−^ yki^OE^* clones and development of the host larvae. Surprisingly, the levels of HSF in cells in *lgl^−^ yki^OE^* clones (N=38) were similar to those in the surrounding *lgl^−^/lgl^+^* cells (**Fig. e-e’**); this was confirmed by examination of intensity profiles for GFP and HSF signals which showed that the general intensity for HSF was similar in GFP-positive clonal and adjacent GFP-negative cells (**Fig. 9h**). Co-staining for Lamin C of wing discs carrying *lgl^−^ yki^OE^* clones (**Fig. 9e’**) further showed that the HSF remained similarly enriched in nuclei in the clonal mutant and the surrounding normal cells.

Down-(**Fig. 9f-P**) or up-regulation (**Fig. 9g-g’**) of HSF in *lgl^−^ yki^OE^ HSF^RNAi^* or *lgl^−^ yki^OE^ HSF^OE^* clones, respectively, was confirmed by immunostaining with HSF-antibody. Although the HSF in *HSF^OE^* transgene was tagged with GFP, we used HSF-immunostaining to clearly demarcate the HSF expression from the GFP fluorescence seen in MARCM clonal cells. Near-complete absence of HSF in *lgl^−^yki^OE^ HSF^RNAi^* clones affected neither their frequency, nor size (**Fig. 9j**), and nor survival of the host larvae (**Fig. 9k**). Unlike in *lgl^−^* clones, HSF over-expression in *lgl^−^ yki^OE^ HSF^OE^* clones affected neither the growth of clones nor the survival of hosts (data not shown). These results thus indicate that while over-expression of HSF provides competitive advantage to *lgl^−^* clones, it is not required for survival and establishment of tumorigenic *lgl^−^ yki^OE^* clones.

## Discussion

The availability of sophisticated targeting techniques like the GAL4/UAS binary system for conditional expression and the FLP-FRT based clonal analysis that permit up- or down-regulation of the desired gene/s in specific tissues or groups of cells at defined time of development (Gangwani et al., 2020) have made *Drosophila* a convenient and informative model for studying the signaling pathways controlling tumor progression, which are largely conserved between *Drosophila* and humans (Ugur et al., 2016; Kanda and Igaki, 2020; Mirzoyan et al., 2019). The presence of mutant cells juxtaposed with wild-type cells in the somatic clone carrying mosaic discs allows knowing the specific time when a cell started on its tumorigenic journey and also permits a study of effects of the host tissue’s microenvironment (Miles et al., 2011; Ohsawa, 2019; Kanda and Igaki, 2020) that affects survival of the potential neoplastic cell.

Elevation in levels of different Hsps has been widely reported in mammalian tumors (Workman, 2004; Basto et al., 2007; Aswad and Liu, 2021; Calderwood and Murshid, 2017; Chatterjee and Burns, 2017; Cyran and Zhitkovich, 2022). However, their expression and role in relation to the initial phase of tumor growth has not been examined earlier. Taking advantage of the unique feature of clonal analysis in *Drosophila,* we show for the first time that the various constitutively expressed Hsps, but not the stress-inducible Hsp70, become more abundant in growing Lgl loss and Yorkie gain associated tumors since their early stages, even before they begin to be transformed as reflected by the absence of F-actin accumulation (Khan et al., 2013). The further elevation in levels of the constitutively expressed Hsps in aggressively growing tumors agrees with and supplements earlier reports of enhanced levels of different Hsps in mammalian cancers at advanced stages of tumorigenesis (Lang et al., 2019; Cyran and Zhitkovich, 2022; Chatterjee and Burns, 2017).

Cell competition acts a check point to selectively eliminate unfit/mutated cells from the growing tissue via cell-cell interaction and thus ensures tissue or organ fitness (Nagata and Igaki, 2018). Cell competition, resulting in elimination of oncogenic cells or selection of fitter/stronger cells, is influenced by mutations that cause loss of apicobasal cell polarity and/or affect signaling activity of the Hippo pathway (Brumby and Richardson, 2003; Tyler et al., 2007; Nagata and Igaki, 2018). Hsp90 has anti-apoptotic (Arya et al., 2007) role and also regulates cell-cycle exit in *Drosophila* so that its elevated levels protect the tumor progenitor cells from death inducing signals from surrounding normal cells. Our study reveals Hsp83 to be an important regulator of cell-competition since down-regulation of Hsp83 levels reduced the early survival as well as the later progression of the surviving *lgl^−^ yki^OE^* while its over-expression converted the normally dis-advantaged *lgl^−^* clones (Menéndez et al., 2010) to be competitively successful and survive in larger numbers. *Drosophila* Hsp83, like its mammalian Hsp90 ortholog, is known to regulate autophagy by mediating the stability or/and activity of signaling proteins like Ulk1, Beclin1, Akt and LAMP-2A (Wang et al., 2016) so that its down-regulation in *lgl^−^ yki^OE^* clones makes them sensitive to apoptotic death. In a recent study (Choutka et al., 2017) it has been shown that loss of Hsp83 elevates levels of the effector caspase DCP-1, which can also activate compensatory autophagy. Generally, the compensatory autophagy makes the tumor cells super-competitors by non-autonomously upregulating autophagy in surrounding wild-type cells so that the tumorigenic cells can grow and invade (Nagata and Igaki, 2018; Katheder et al., 2017). It is tempting to speculate that within a *lgl^−^ yki^OE^ Hsp83^RNAi^* clone, a few cells with high DCP-1 may also induce compensatory autophagy in neighboring clonal cells so that the combined effect of apoptosis and compensatory autophagy in different cells in *lgl^−^ yki^OE^ Hsp83^RNAi^* clones together results in a poor survival of the clone as a whole, as seen in our study.

Elevated levels of Hsp83 also appear to be necessary for transformation of the tumorigenic clones at later stages since the F-actin accumulation in the surviving *lgl^−^ yki^OE^ Hsp83^RNAi^* clones at 72hr ACI was perceptibly lower than in similar age *lgl^−^ yki^OE^* or *lgl^−^* clones. Unlike the suppressive effects of down-regulation of Hsp83 on survival and continuance of tumorous clones, its over-expression in *lgl^−^ yki^OE^ Hsp83^OE^* or *lgl^−^ Hsp83^OE^* clones did not have noticeable tumor-enhancing affects in wing discs but the *lgl^−^ Hsp83^OE^* clones appeared generally larger in eye discs with the emerging flies showing a higher incidence of necrotic patches in and around eyes. The different responses of *lgl^−^* clones in wing and eye-antennal discs to further elevated expression of Hsp83 reflects differences in the regulatory networks in these two tissues. It is also possible that *Hsp83^OE^* co-expression in the clones did not elevate levels of Hsp83 much beyond the already elevated levels in *lgl^−^ yki^OE^* or *lgl^−^* tumorous clones in wing discs and thus had little additive effect in this tissue.

Like Hsp83, Hsp60 and Hsp27 levels were also found to be elevated in clones and whole disc tumors. *Drosophila melanogaster* has four different Hsp60 proteins, named Hsp60A to Hsp60D (Sarkar and Lakhotia, 2005; Sarkar et al., 2006). The anti-Hsp60 antibody used in this study appears to identify all the four Hsp60 members. However, our other studies (Singh and Lakhotia, in preparation) suggest that the Hsp60D plays an important role in the progression of *lgl^−^ yki^OE^* clones since down regulation of the Hsp60D suppressed growth of *lgl^−^ yki^OE^* clones.

Expression of the Hsp70 family proteins in tumor clones in our study was found to follow a unique course since while the constitutively expressed Hsc70 family proteins exhibited a general increase similar to the other constitutively expressed Hsps, the stress-inducible Hsp70 was found to be expressed only at a later stage (72hr or more ACI) of *lgl^−^ yki^OE^* clone growth when most of them already had high F-actin aggregation and thus may have been in the process of getting transformed. More interestingly, only a small number of cells within the tumor exhibited the much higher levels of Hsp70. A comparable pattern of expression of stress-inducible Hsp70 has so far not been reported in any other study on tumor cells although some human tumors do not show significantly elevated level of the inducible Hsp70 (Cyran and Zhitkovich, 2022). A more detailed account of this intriguing observation will be presented elsewhere (Singh and Lakhotia, in preparation).

Activation of heat shock genes following cell stress is majorly regulated by the HSF, which unlike in mammals is encoded by a single gene in *Drosophila.* Many studies on mammalian cancers have suggested the HSF1 to play an important role in modulation of Hsp levels in tumor cells and cancer progression (Alasady and Mendillo, 2020; Dai et al., 2007; Dai and Sampson, 2016; Cyran and Zhitkovich, 2022). In contrast, our results show that while down-regulation of Hsp83 suppressed survival and growth of the *lgl^−^* or *lgl^−^ yki^OE^* clones and consequently improved survival of the clone-carrying larvae, down-regulation of HSF in the clones affected neither their survival and growth nor the host’s survival. This observation indicates that the elevation of constitutively expressed members of Hsp families in *Drosophila lgl^−^ yki^OE^* tumors is not dependent upon HSF. This is also borne by our finding that the HSF levels were not elevated in these tumorous clones. The developmental expression patterns of Hsps like Hsp83, Hsp60, sHsps are regulated by diverse regulatory pathways independent of HSF. For example, mammalian Hsp90 is also regulated by NF-kappa B. nuclear factor interleukin-6 (NF-IL6), and by signal transducer and activator of transcription-3 (STAT-3) signalling pathways under different conditions (Stephanou et al., 1998; Ammirante et al., 2008). Interestingly, *Drosophila hsp83* gene also shows binding sites for Relish, the fly homolog of NF-kappa B (Copley et al., 2007). Apparently, like the HSF-independent developmental expression of constitutively expressed Hsps (Cohen and Meselson, 1985; Hoffman and Corces, 1986; Hultmark et al., 1986; Xiao and Lis, 1989), their elevated expression in *Drosophila* tumorous clones examined in this study may also be dependent upon signalling pathways other than HSF. This needs further analysis.

The observed better survival and transformation of *lgl^−^* clones following over-expression of HSF contrasts with that in *lgl^−^ yki^OE^* clones. In our other study (Singh and Lakhotia, under preparation) we found that *lgl^−^ Ras^V12^* clones need HSF for Hsp70 expression, and over-expression of HSF makes them grow more aggressively. Likewise, HSF-overexpression indicates poor prognosis for many human cancers (Dai et al., 2007; Cyran and Zhitkovich, 2022; Meng et al., 2010). Thus, HSF’s role in tumorigenesis may be context-dependent.

The various Hsps work in close association with their respective co-chaperones (Altinok et al., 2021; Joshi et al., 2018). Expression patterns of the diverse co-chaperones and their roles in survival and progression of the tumorous clones remain to be examined. Future studies in this direction using the fly model of clonal analysis of tumor progression will indeed be very useful.

Different Hsps play fundamental roles in processes of signal transduction, cell cycle progression and apoptosis, cell proliferation and survival, all of which contribute to their involvement, via their client proteins and co-chaperones, in invasion, metastasis and tumor angiogenesis (Lang et al., 2019; Joshi et al., 2018). Our experimental strategy has revealed some results that contradict the generally believed roles of different Hsps in progression of cancer. We believe that this experimental model will be very useful in unravelling the roles of different Hsps, their cochaperones and HSF during different stages of tumor growth more precisely. Since the Hsps have gained interest as promising anticancer drug targets (Calderwood and Murshid, 2017; Chatterjee and Burns, 2017; Basto et al., 2007; Aswad and Liu, 2021; Lang et al., 2019), a better understanding of specific roles of different Hsps and HSF in tumor initiation and establishment will help in better therapeutic applications.

## Acknowledgements

Gunjan Singh was supported by the University Grants Commission (New Delhi). SCL acknowledges SERB (Govt. of India) Distinguished Fellowship. We thank Prof. Pradip Sinha (IIT, Kanpur, India), and the Bloomington Drosophila Stock Centre for different fly stocks, Prof. M. Evgen’ev (Russia) and Prof. Robert Tanguay (Canada) for providing Hsp70 and Hsp83 antibodies, respectively and Prof. J. T. Lis for the HSF-overexpression stock and HSF antibody. We also acknowledge the kind gift of Anti-Hsp/Hsc70 antibody by late Prof. S. Lindquist.

## References

Agrawal N, Joshi S, Kango M et al. (1995). Epithelial hyperplasia of imaginal discs induced by mutations in Drosophila tumor suppressor genes: growth and pattern formation in genetic mosaics. Dev Biol, 169(2), 387–98. doi:10.1006/dbio.1995.1155.

Alasady MJ, Mendillo ML (2020). The Multifaceted Role of HSF1 in Tumorigenesis. Adv Exp Med Biol, 1243, 69–85. doi: 10.1007/978-3-030-40204-4_5.

Altinok S, Sanchez-Hodge R, Stewart M et al. (2021). With or without you: Co-chaperones mediate health and disease by modifying chaperone function and protein triage. Cells, 10(11), 3121. doi:doi:10.3390/cells10113121.

Ammirante M, Rosati A, Gentilella A et al. (2008). The activity of hsp90 alpha promoter is regulated by NF-kappa B transcription factors. Oncogene, 27(8), 1175–8. doi:10.1038/sj.onc.1210716.

Arya R, Mallik M, Lakhotia SC (2007). Heat shock genes – integrating cell survival and death. J. Biosciences, 32(3), 595–610. doi:10.1007/s12038-007-0059-3.

Aswad A, Liu T (2021). Targeting heat shock protein 90 for anti-cancer drug development. Adv Cancer Res, 152, 179–204. doi:10.1016/bs.acr.2021.03.006.

Baker NE (2020). Emerging mechanisms of cell competition. Nature Reviews Genetics, 21(11), 683–697. doi:10.1038/s41576-020-0262-8.

Basto R, Gergely F, Draviam VM et al. (2007). Hsp90 is required to localise cyclin B and Msps/ch-TOG to the mitotic spindle in Drosophila and humans. J Cell Sci, 120(Pt 7), 1278–87. doi:10.1242/jcs.000604.

Birch-Machin I, Gao S, Huen D et al. (2005). Genomic analysis of heat-shock factor targets in Drosophila. Genome Biol, 6(7), R63. doi:10.1186/gb-2005-6-7-r63.

Brumby AM, Richardson HE (2003). scribble mutants cooperate with oncogenic Ras or Notch to cause neoplastic overgrowth in Drosophila. Embo J, 22(21), 5769–79. doi:10.1093/emboj/cdg548.

Calderwood SK (Ed.) (2007). Cell stress proteins. (Berlin: Springer)

Calderwood SK, Gong J (2016). Heat shock proteins promote cancer: It’s a protection eacket. Trends in Biochemical Sciences, 41(4), 311–323. doi:https://doi.org/10.1016/j.tibs.2016.01.003.

Calderwood SK, Murshid A (2017). Molecular chaperone accumulation in cancer and decrease in Alzheimer’s disease: The potentialroles of HSF1. Frontiers in Neuroscience, 11. doi:10.3389/fnins.2017.00192.

Capdevila J, Estrada MP, Sánchez-Herrero E et al. (1994). The Drosophila segment polarity gene patched interacts with decapentaplegic in wing development. Embo Journal, 13(1), 71–82.

Capdevila J, Guerrero I (1994). Targeted expression of the signaling molecule decapentaplegic induces pattern duplications and growth alterations in Drosophila wings. EMBO Journal, 13(19), 4459–4468.

Chatterjee S, Burns TF (2017). Targeting heat shock proteins in cancer: A promising therapeutic approach. International Journal of Molecular Sciences, 18(9), 1978.

Choutka C, DeVorkin L, Go NE et al. (2017). Hsp83 loss suppresses proteasomal activity resulting in an upregulation of caspase-dependent compensatory autophagy. Autophagy, 13(9), 1573–1589. doi:10.1080/15548627.2017.1339004.

Cohen RS, Meselson M (1985). Separate regulatory elements for the heat-inducible and ovarian expression of the Drosophila hsp26 gene. Cell, 43(3 Pt 2), 737–46. doi:10.1016/0092-8674(85)90247-8.

Copley RR, Totrov M, Linnell J et al. (2007). Functional conservation of Rel binding sites in drosophilid genomes. Genome research, 17(9), 1327–1335.

Cyran AM, Zhitkovich A (2022). Heat shock proteins and HSF1 in cancer. Front. Oncology, 12, 860320. doi:10.3389/fonc.2022.860320.

Dai C, Sampson SB (2016). HSF1: Guardian of proteostasis in cancer. Trends in Cell Biology, 26(1), 17–28. doi: doi.org/10.1016/j.tcb.2015.10.011.

Dai C, Whitesell L, Rogers AB et al. (2007). Heat shock factor 1 is a powerful multifaceted modifier of carcinogenesis. Cell, 130(6), 1005–18. doi:10.1016/j.cell.2007.07.020.

Elefant F, Palter KB (1999). Tissue-specific expression of dominant negative mutant Drosophila HSC70 causes developmental defects and lethality. Mol Biol Cell, 10(7), 2101–17. doi:10.1091/mbc.10.7.2101.

Feder ME, Hofmann GE (1999). Heat-shock proteins, molecular chaperones, and the stress response: evolutionary and ecological physiology. Annu Rev Physiol, 61, 243–82. doi:10.1146/annurev.physiol.61.1.243.

Gangwani K, Snigdha K, Atkins M et al. (2020). Drosophila cancer modeling using the eye imaginal discs. In Singh A & Kango-Singh M (Eds.), Molecular Genetics of Axial Patterning, Growth and Disease in Drosophila Eye (pp. 259–291). Switzerland: Springer Nature

Hanahan D, Weinberg RA (2000). The hallmarks of cancer. Cell, 100(1), 57–70. doi:10.1016/s0092-8674(00)81683-9.

Hanahan D, Weinberg RA (2011). Hallmarks of cancer: the next generation. Cell, 144(5), 646–74. doi:10.1016/j.cell.2011.02.013.

Hoffman E, Corces V (1986). Sequences involved in temperature and ecdysterone-induced transcription are located in separate regions of a Drosophila melanogaster heat shock gene. Mol Cell Biol, 6(2), 663–73. doi:10.1128/mcb.6.2.663-673.1986.

Hultmark D, Klemenz R, Gehring WJ (1986). Translational and transcriptional control elements in the untranslated leader of the heat-shock gene hsp22. Cell, 44(3), 429–38. doi:10.1016/0092-8674(86)90464-2.

Joshi S, Wang T, Araujo TLS et al. (2018). Adapting to stress — chaperome networks in cancer. Nature Reviews Cancer, 18(9), 562–575. doi:10.1038/s41568-018-0020-9.

Kanda H, Igaki T (2020). Mechanism of tumor-suppressive cell competition in flies. Cancer Sci, 111(10), 3409–3415. doi:10.1111/cas.14575.

Katheder NS, Khezri R, O’Farrell F et al. (2017). Microenvironmental autophagy promotes tumour growth. Nature, 541(7637), 417–420. doi:10.1038/nature20815.

Khan S, Bajpai A, Alam M et al. (2013). Epithelial neoplasia in Drosophila entails switch to primitive cell states. Proceedings of the National Academy of Sciences of the United States of America, 110(24), 72. doi:10.1073/pnas.1212513110.

Lang BJ, Guerrero-Giménez ME, Prince TL et al. (2019). Heat shock proteins are essential components in transformation and tumor progression: Cancer cell intrinsic pathways and beyond. Int J Mol Sci, 20(18). doi:10.3390/ijms20184507.

Lee T, Luo L (1999). Mosaic analysis with a repressible cell marker for studies of gene function in neuronal morphogenesis. Neuron, 22(3), 451–61. doi:10.1016/s0896-6273(00)80701-1.

Lee T, Luo L (2001). Mosaic analysis with a repressible cell marker (MARCM) for Drosophila neural development. Trends Neurosci, 24(5), 251–4. doi:10.1016/s0166-2236(00)01791-4.

Lindquist S, Craig EA (1988). The heat-shock proteins. Annu Rev Genet, 22, 631–77. doi:10.1146/annurev.ge.22.120188.003215.

Menéndez J, Pérez-Garijo A, Calleja M et al. (2010). A tumor-suppressing mechanism in Drosophila involving cell competition and the Hippo pathway. Proc Natl Acad Sci U S A, 107(33), 14651–6. doi:10.1073/pnas.1009376107.

Meng L, Gabai VL, Sherman MY (2010). Heat-shock transcription factor HSF1 has a critical role in human epidermal growth factor receptor-2-induced cellular transformation and tumorigenesis. Oncogene, 29(37), 5204–5213. doi:10.1038/onc.2010.277.

Miles WO, Dyson NJ, Walker JA (2011). Modeling tumor invasion and metastasis in Drosophila. Disease Models and Mechanisms, 4(6), 753–761.

Millburn GH, Crosby MA, Gramates LS et al. (2016). FlyBase portals to human disease research using Drosophila models. Dis Model Mech, 9(3), 245–52. doi:10.1242/dmm.023317.

Mirzoyan Z, Sollazzo M, Allocca M et al. (2019). Drosophila melanogaster: A Model Organism to Study Cancer. Front Genet, 10, 51. doi:10.3389/fgene.2019.00051.

Nagata R, Igaki T (2018). Cell competition: Emerging mechanisms to eliminate neighbors. Development, Growth & Differentiation, 60(9), 522–530. doi:https://doi.org/10.1111/dgd.12575.

Ohsawa S (2019). Elimination of oncogenic cells that regulate epithelial homeostasis in Drosophila. Dev Growth Differ, 61(5), 337–342. doi:10.1111/dgd.12604.

Ray M, Singh G, Lakhotia SC (2019). Altered levels of hsromega lncRNAs further enhance Ras signaling during ectopically activated Ras induced R7 differentiation in Drosophila. Gene Expr Patterns, 33, 20–36. doi:10.1016/j.gep.2019.05.002.

Richter K, Haslbeck M, Buchner J (2010). The heat shock response: life on the verge of death. Mol Cell, 40(2), 253–66. doi:10.1016/j.molcel.2010.10.006.

Sarkar S, Arya R, Lakhotia SC (2006). Chaperonins: in life and death. In Sreedhar ASSrinivas UK (Eds.), Stress responses: A molecular biology approach (pp. 43–60). Trivandrum, India: Signpost.

Sarkar S, Lakhotia SC (2005). The Hsp60C gene in the 25F cytogenetic region in Drosophila melanogaster is essential for tracheal development and fertility. J Genet, 84(3), 265–81. doi:10.1007/bf02715797.

Stephanou A, Isenberg DA, Akira S et al. (1998). The nuclear factor interleukin-6 (NF-IL6) and signal transducer and activator of transcription-3 (STAT-3) signalling pathways co-operate to mediate the activation of the hsp90beta gene by interleukin-6 but have opposite effects on its inducibility by heat shock. Biochem J, 330(Pt 1)(Pt 1), 189–95. doi:10.1042/bj3300189.

Tyler DM, Li W, Zhuo N et al. (2007). Genes affecting cell competition in Drosophila. Genetics, 175(2), 643–57. doi:10.1534/genetics.106.061929.

Ugur B, Chen K, Bellen HJ (2016). Drosophila tools and assays for the study of human diseases. Dis Model Mech, 9(3), 235–44. doi:10.1242/dmm.023762.

Velazquez JM, Lindquist S (1984). hsp70: nuclear concentration during environmental stress and cytoplasmic storage during recovery. Cell, 36(3), 655–62. doi:10.1016/0092-8674(84)90345-3.

Vogelstein B, Fearon ER, Hamilton SR et al. (1988). Genetic alterations during colorectal-tumor development. N Engl J Med, 319(9), 525–32. doi:10.1056/nejm198809013190901.

Wang B, Chen Z, Yu F et al. (2016). Hsp90 regulates autophagy and plays a role in cancer therapy. Tumour Biol, 37(1), 1–6. doi:10.1007/s13277-015-4142-3.

Workman P (2004). Altered states: selectively drugging the Hsp90 cancer chaperone. Trends Mol Med, 10(2), 47–51. doi:10.1016/j.molmed.2003.12.005.

Xiao H, Lis JT (1989). Heat shock and developmental regulation of the Drosophila melanogaster hsp83 gene. Mol Cell Biol, 9(4), 1746–53. doi:10.1128/mcb.9.4.1746-1753.1989.

Yepiskoposyan H, Egli D, Fergestad T et al. (2006). Transcriptome response to heavy metal stress in Drosophila reveals a new zinc transporter that confers resistance to zinc. Nucleic Acids Res, 34(17), 4866–77. doi:10.1093/nar/gkl606.

Yokota J (2000). Tumor progression and metastasis. Carcinogenesis, 21(3), 497–503. doi:10.1093/carcin/21.3.497.

Zaarur N, Gabai VL, Porco JA, Jr. et al. (2006). Targeting heat shock response to sensitize cancer cells to proteasome and Hsp90 iInhibitors. Cancer Research, 66(3), 1783–1791. doi:10.1158/0008-5472.Can-05-3692.

